# Molecular pathway analysis indicates a distinct metabolic and immune phenotype in women with right-sided colon cancer

**DOI:** 10.1101/659151

**Authors:** Yazhi Sun, Varvara Mironova, Ying Chen, Elliott P.F. Lundh, Qian Zhang, Yuping Cai, Vasilis Vasiliou, Yawei Zhang, Rolando Garcia-Milian, Sajid A. Khan, Caroline H. Johnson

## Abstract

Colon cancer is the third most commonly diagnosed cancer in the United States. Recent reports have shown that the location of the primary tumor is of clinical importance. Patients with right-sided cancers (RCCs) (tumors arising between the cecum and proximal transverse colon) have poorer clinical outcomes than those with left-sided colon cancers (LCCs) (tumors arising between the distal transverse colon and sigmoid colon, excluding the rectum). Interestingly, women have a lower incidence of colon cancer than men do. However, women have a higher propensity for RCC than men. Identification of gene expression differences between RCC and LCC is considered a potential means of prognostication. Furthermore, studying colon cancer sidedness could reveal important predictive markers for response to various treatments. This study provides a comprehensive bioinformatic analysis of various genes and molecular pathways that correlated with sex and anatomical location of colon cancer using four publicly available annotated datasets housed in the National Center for Biotechnology Information’s Gene Expression Omnibus (GEO). We identified differentially expressed genes in tumor tissues from women with RCC, which showed attenuated energy and nutrient metabolism when compared to women with LCC. Specifically, we showed that the downregulation of 5’ AMP-activated protein kinase alpha subunit (*AMPKα*) and downregulated anti-tumor immune response in women with RCC. This difference was not seen when comparing tumor tissues from men with RCC to men with LCC. Therefore, women with RCC may have a specific metabolic and immune phenotype which accounts for differences in prognosis and treatment response.

## Introduction

Colon cancer is the third most commonly diagnosed cancer, and the second leading cause of cancer-related death in the United States (Aran et al., 2016; Siegel et al., 2017). The incidence and mortality rate of colon cancer has been steadily declining for past several years in Western countries. This is primarily due to advances in screening programs and lifestyle changes (Haggar and Boushey, 2009; Waly and Ali, 2018). Colonoscopy and fecal-occult blood tests are the most commonly used screening tools that have shown a significant reduction in incidence and mortality. However, the incidence of colon cancer continues to increase in countries that are transitioning into high-income economies, such as Eastern Asia countries and Eastern European countries. This is possibly due to adoption of Westernized diets that are high in fat and various environmental exposures (Larsson et al., 2005; Haggar and Boushey, 2009). Despite the decrease in incidence rate in Western countries and the development of techniques that have improved diagnosis and treatment of colon cancer in recent years, mortality is still high (14.8 per 100,000 person). Worldwide mortality rate is approximately half that of the incidence rate (Haggar and Boushey, 2009; Lee et al., 2017b).

Recent reports have shown that the location of the primary tumor is of clinical importance (Gervaz et al., 2016). The left and right side of the colon have distinct embryologic origins, vasculature, and differing gene expression patterns (Gervaz et al., 2004). Furthermore, the two sides of the colon have different exposures to environmental compounds, microbiome density, and metabolite distribution (Lee et al., 2017a). Cancer stemming from these two regions are known to exhibit different epidemiological, histological and clinical characteristics (Lee et al., 2017b). For instance, patients with right-sided colon cancers (RCCs) (tumors arising between the cecum and proximal transverse colon) are more likely to be women, of more advanced age, and have worse clinical outcomes compared to those with left-sided colon cancers (LCCs) (tumors arising between the distal transverse colon and sigmoid colon, excluding the rectum) (Benedix et al., 2010; Lee et al., 2017b). Therefore, the pathophysiology that control RCC versus LCC are likely different, but, at present, not well characterized. Studies that examine location of colon cancer, particularly in women with RCC, are needed to improve existing preventative and therapeutic options for these patients.

Bioinformatics approaches for analyzing genome-wide transcriptomic data can assess the relationship between gene expression and causal mechanisms, and are enabling interpretation of these high-dimensional datasets. Enrichment analysis, for example, evaluates high-dimensional data at the level of gene sets, and provides a large-scale comparison at the molecular pathway and disease process level instead of examining individual genes (Subramanian et al., 2005). In a recent paper, an enrichment analyses of gene expression correlation between RCC and LCC patients was carried out using the GSE14333 dataset from the public database National Center for Biotechnology Information Gene Expression Omnibus (GEO) (Peng et al., 2018). The study revealed molecular pathways that were differentially correlated with tumor development in these two regions of the colon. However, in this study, only one population cohort was examined and did not correct for multiple comparison testing (Peng et al., 2018). Another study used data from both the Cancer Genome Atlas (TCGA) and GSE14333 to examine somatic mutations, genome-wide mRNA and miRNA, and DNA methylation profiles associated with RCC and found a correlation in the phosphoinositide 3-kinase (PI3K) signaling pathway (Hu et al., 2018).

In the present study, we retrieved gene expression profiles of patients with colon cancer from four GEO datasets to identify significant gene expression differences and their reproducibility between men and women with RCC and LCC. We identified groups of related genes residing in one or multiple molecular pathways that were commonly altered in women and men with RCC or LCC using enrichment analysis, and showed reproducibility of results between the datasets. Thus, we identified molecular differences between primary colon tumor location in men and women, and generated hypotheses pertaining to the causal mechanisms for clinical and epidemiological differences between these groups of samples.

## Materials and Methods

### Data collection

Gene expression profiles were retrieved from the public database GEO, including microarray datasets GSE41258, GSE39582, GSE37892, and GSE14333. GSE41258 was generated using the GPL96 platform for transcriptome analysis (Affymetrix Human Genome U133A array), while the other datasets were generated using the GPL570 platform (Affymetrix Human Genome U133Plus 2.0 arrays). The GPL570 platform is an updated version of GPL96, with the addition of 6,500 genes. All datasets were downloaded from the GEO database in Qlucore Omics Explorer (Version 3.3, Qlucore AB, Lund, Sweden).

### Sample selection and inclusion criteria

To stratify samples by anatomical location, we defined RCC cases using the ontology terms “right”, “ascending”, “cecum”, “hepatic flexure”. For LCC terms “distal”, “descending”, “sigmoid”, “splenic flexure” were used. Only primary tumor samples were selected including those recorded as “carcinoma” and “adenocarcinoma”. Furthermore, only those samples annotated with information regarding patient sex were considered. Samples that fell out of these strict inclusion criteria bounds were excluded from the study. The four selected datasets had more than 100 samples each remaining after application of the inclusion/exclusion criteria (Table 1). Qlucore Omics Explorer was used for data selection and categorizing (Version 3.3, Qlucore AB, Lund, Sweden).

**Table 1.** Patient characteristics from each dataset.

|  | GSE39582 | GSE37892 | GSE14333 | GSE41258 |
| --- | --- | --- | --- | --- |
| **LCC Total (Number of patients)** | 75 | 72 | 122 | 75 |
| Women (Number (% of LCC total)) | 31 (41.3) | 34 (47.2) | 45 (38.9) | 37 (49.3) |
| Men (Number (% of LCC total)) | 44 (58.7) | 38 (52.8) | 77 (63.1) | 38 (50.7) |
| **RCC Total (Number of patients)** | 48 | 57 | 125 | 62 |
| Women (Number (% of RCC total)) | 19 (39.6) | 26 (45.6) | 66 (52.8) | 30 (48.4) |
| Men (Number (% of RCC total)) | 29 (60.4) | 31 (54.4) | 59 (47.2) | 32 (51.6) |

### Analysis of patient characteristics

Selected sample annotation files were exported from Qlucore Omics Explorer and analyzed in SAS software (SAS Institute Inc., Cary, NC, USA). The mean age of patients was compared to sex and cancer anatomical location (student’s t-test, p<0.05 statistically significant). We also used Ingenuity Pathway Analysis (IPA) for additional pathway analysis between women with RCC and LCC.

### Identification of differentially expressed genes

All gene expression data was log-transformed in Qlucore Omics Explorer to stabilize the variance, compress the range of data, and normalize the distribution of the data. Student’s t-test was used comparing women with RCC as the selected group to women with LCC to obtain differentially expressed genes specific to cancer location. We also compared women with RCC to men with RCC to investigate the influence of sex on gene expression. Results were thus differentially expressed with respect to women with RCC. Then we calculated adjusted p-values (Benjamini-Hochberg False Discovery Rate) to account for multiple comparisons, and log2 transformed fold change for each gene.

### Pathway analysis and comparison for reproducibility

Pathway analysis was conducted using MetaCore™ software (GeneGo, San Diego, CA) which is a systems biology analysis suite to identify altered gene functions and pathways. The Affymetrix Human Genome U133A Array was used for probe set annotations. The differentially expressed gene lists with adjusted p-value and log2 transformed were uploaded with a threshold of q-value < 0.1. The results of the enrichment analysis provided pathway maps, which were determined to be of statistical significance using a FDR < 0.05. We also used Ingenuity Pathway Analysis (IPA, QIAGEN Redwood City, USA) to look for additional pathways as MetaCore™ and IPA have different knowledge bases and may reveal possible differences in regulation of different signaling pathways related to colon cancer.

IPA network analysis was used to map differentially expressed genes between women with RCC and LCC. Differentially expressed genes, which interact with other molecules in the Ingenuity Knowledge Base, are identified as network-eligible molecules. These serve as “seeds” for generating networks (green are down regulated; red are upregulated) through the IPA network generating algorithm. Network-eligible molecules are combined into networks that maximize their interconnectedness with each other relative to all molecules they are connected to in the Ingenuity Knowledge Base. Generated networks are scored based on the probability of finding observed number of network-eligible molecules in a given network by random chance.

## Results

### Patient characteristics

For each publicly available dataset obtained from GEO, information regarding platform type, sample size and patient characteristics can be observed on Table 1. The mean age of all patients with RCC was higher than all patients with LCC across all datasets but only reached statistical significance in GSE14333 (p-value < 0.001), see Table 2. This is in accordance with previous studies (Gonzalez et al., 2001; Benedix et al., 2010). The mean age of women with all cancer locations compared to men was also only statistically significantly higher in GSE14333 (p-value 0.006).

**Table 2.**
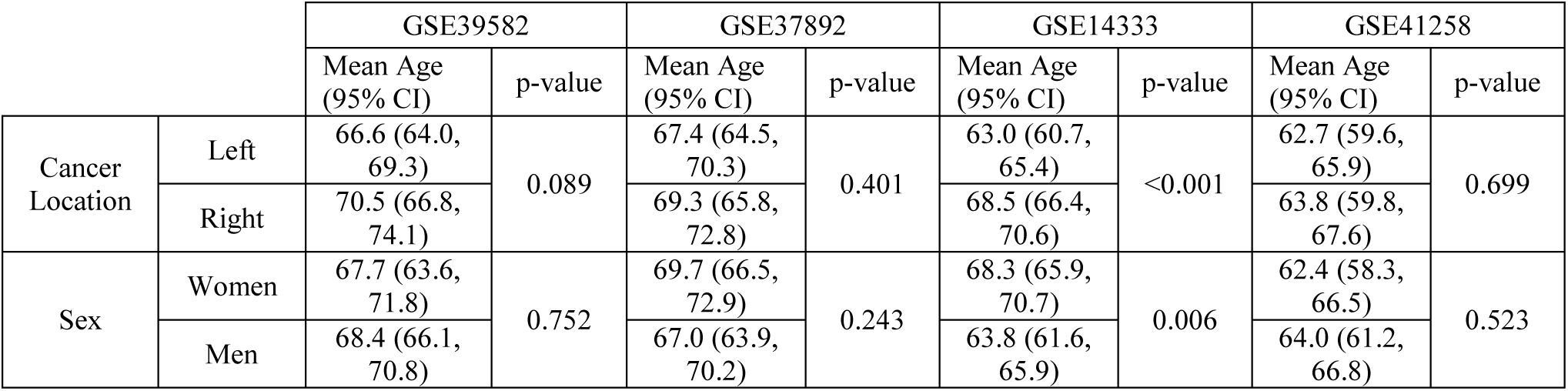
Mean age of patients compared by sex and cancer anatomical location in each dataset (t-test, p<0.05 statistically significant).

### Principal component analysis (PCA) of all differentially expressed genes across the group comparisons

We carried out PCA analysis of gene expression profiles from tumor tissues taken from women with RCC. These profiles showed some separation from women with LCC and from men with RCC in the GSE41258 dataset (Figures 1A and 1B). Figure 1A shows the PCA scores plot of data from GSE41258 comparing gene expression values from women with RCC to women with LCC, q-value < 0.1. Similarly, Figure 1B shows a PCA comparing women with RCC and men with RCC (q-value < 0.1 by student’s t-test). Overall, the PCA models revealed that there are differences in gene expression between the sample groups, however the maximal difference appears to be between women and men with RCC compared to women with RCC and LCC.

**Figure 1A.**
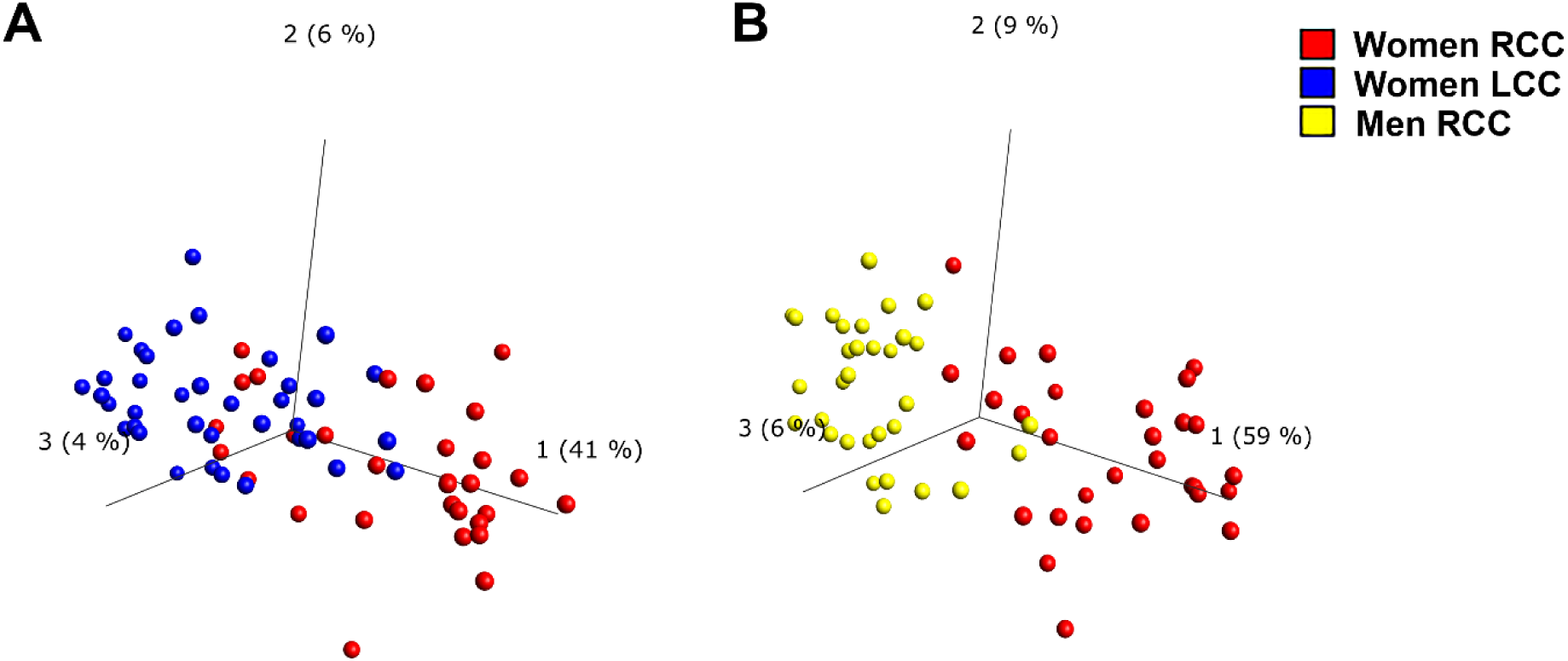
PCA plot of data from GSE41258 comparing gene expression values from women with RCC to women with LCC, q-value < 0.1. Principal component 1 explains the genes that contribute to 41% of the total variance between the two sets of samples. **1B**. PCA plot of GSE41258 when comparing women with RCC and men with RCC (q-value < 0.1 by student’s t-test). Principal component 1 explains the genes that contribute to 56% of the total variance between the two patient groups. Supplemental Figure 1 shows PCA scores plots for the other four datasets.

### Identification of differently expressed genes between RCCs and LCCs in women and men

Three datasets (GSE39582, GSE14333 and GSE41258) were observed to have statistically significant differences in gene expression between tumors in women with RCC and LCC after correction for multiple comparisons. Table 3 shows the genes that are reproducibly dysregulated in ≥ 2 datasets. Many of the genes had fold changes of ≥ 2 (log2FC ≥ 1 or ≤ −1) when comparing women with RCC to women with LCC. These included AT-Rich Interaction Domain 3A (*ARID3A*), Special AT-Rich Sequence-Binding Protein 2 (*SATB2*), and Troponin C2 Fast Skeletal Type (*TNNC2*), which were downregulated in women with RCC. Conversely, homeobox C6 (*HOXC6*) was upregulated in women with RCC.

**Table 3.** Differential expression of genes in colon tumor from women with RCC compared to women with LCC that reproducibly occur in two or more datasets. Log2 fold changes displayed are with respect to women with RCC. *Also, differentially regulated in men with RCC compared to men with LCC.

|  | GSE14333 |  | GSE41258 |  | GSE39582 |  |
| --- | --- | --- | --- | --- | --- | --- |
| Gene Symbol | q-value | Log2 fold change | q-value | Log2 fold change | q-value | Log2 fold change |
| ACOT8 |  |  | 7.88E-02 | -0.36 | 3.41E-02 | -0.09 |
| ACSF2 |  |  | 3.06E-02 | -0.76 | 6.77E-02 | -0.06 |
| ACSL6 |  |  | 7.88E-02 | -0.34 | 3.47E-02 | -0.18 |
| ARFGEF2 |  |  | 4.77E-02 | -0.42 | 8.32E-02 | -0.06 |
| ARID3A |  |  | 4.77E-02 | -1.06 | 3.35E-02 | -0.12 |
| ASXL1 |  |  | 4.77E-02 | -0.34 | 4.65E-02 | -0.06 |
| CASP6 |  |  | 9.15E-02 | -0.36 | 7.55E-02 | -0.03 |
| CBFA2T2 |  |  | 9.39E-02 | -0.34 | 1.00E-01 | -0.06 |
| CD24 |  |  | 7.88E-02 | -0.67 | 6.15E-02 | -0.04 |
| GGH |  |  | 9.29E-02 | -0.74 | 2.25E-02 | -0.07 |
| HOXB13 | 2.17E-02 | -0.34 |  |  | 9.62E-02 | -0.10 |
| HOXC6* | 2.73E-03 | 0.18 | 9.88E-02 | 1.02 | 1.10E-02 | 0.30 |
| KIF3B |  |  | 8.07E-02 | -0.43 | 7.51E-02 | -0.04 |
| MUC12* | 5.83E-02 | -0.34 |  |  | 4.65E-02 | -0.18 |
| NEU1 |  |  | 3.06E-02 | -0.62 | 4.25E-02 | -0.06 |
| PDE3A | 4.43E-02 | -0.15 |  |  | 6.51E-02 | -0.10 |
| PFDN4 |  |  | 9.48E-02 | -0.38 | 3.90E-02 | -0.04 |
| PLAGL2 |  |  | 4.77E-02 | -0.60 | 2.25E-02 | -0.09 |
| PNPLA3 |  |  | 9.15E-02 | 0.44 | 3.35E-02 | 0.11 |
| POFUT1 |  |  | 7.88E-02 | -0.67 | 4.25E-02 | -0.07 |
| PRAC1* | 9.26E-13 | -0.56 |  |  | 2.35E-04 | -0.49 |
| RNF43 |  |  | 9.83E-02 | -0.76 | 8.21E-02 | -0.06 |
| SATB2 |  |  | 4.77E-02 | -1.32 | 8.31E-02 | -0.18 |
| STAU1 |  |  | 9.29E-02 | -0.30 | 6.49E-02 | -0.03 |
| TGIF2 |  |  | 9.48E-02 | -0.38 | 2.68E-03 | -0.12 |
| TLE2 |  |  | 9.29E-02 | -0.64 | 2.44E-02 | -0.12 |
| TNNC2 |  |  | 5.93E-02 | -1.89 | 2.78E-02 | -0.10 |
| TSPAN6 |  |  | 9.39E-02 | -0.42 | 1.57E-03 | -0.07 |
| TTI1 |  |  | 4.77E-02 | -0.49 | 6.14E-02 | -0.06 |
| VPS53 | 9.56E-02 | 0.08 |  |  | 7.27E-02 | 0.04 |

To determine whether the differences in gene expression between RCCs and LCCs were sex-specific, we compared gene expression profiles in men with RCC and LCC (Table 4). Again, gene expression data from GSE39582, GSE14333 and GSE41258 revealed differences that were statistically significant, although the number of genes identified was reduced from the comparison of woman with RCC to women with LCC. Of note, *HOXC6* was upregulated, and mucin 12 (*MUC12*) and Prostate Cancer Susceptibility Candidate 1 (*PRAC1*) were downregulated in both men and women with RCC when comparing to LCCs.

**Table 4.** Differential expression of genes in colon tumors from men with RCC compared to men with LCC across three datasets, that reproducibly occur in two or more datasets. Log 2-fold changes displayed are with respect to men with RCC. * Genes that are also differentially regulated in women with RCC when compared to women with LCC.

|  | GSE14333 |  | GSE41258 |  | GSE39582 |  |
| --- | --- | --- | --- | --- | --- | --- |
| Gene Symbol | q-value | Log2 fold change | q-value | Log2 fold change | q-value | Log2 fold change |
| HOXC6* | 5.57E-04 | 0.26 | 3.21E-02 | 1.29 | 1.80E-06 | 0.36 |
| INSL5 | 1.29E-02 | -0.36 |  |  | 5.46E-02 | -0.15 |
| MUC12* | 9.46E-02 | -0.29 |  |  | 2.62E-02 | -0.18 |
| PRAC1* | 3.83E-12 | -0.49 |  |  | 1.61E-03 | -0.42 |
| ZNF345 |  |  | 9.47E-02 | -0.69 | 7.84E-02 | -0.04 |
| ZNF813 |  |  | 9.47E-02 | -0.49 | 2.39E-02 | -0.14 |

Gene expression in tumors from women with RCC was compared to those in tumors from men with RCC, and four datasets were observed to have statistically significant differentially expressed genes after correcting for multiple comparisons (Table 5). Many of the genes had large fold changes of >2, including DEAD-Box Helicase 3, Y-Linked (*DDX3Y*), Lysine Demethylase 5D (*KDM5D*), Ribosomal Protein S4, Y-Linked 1 (*RPS4Y1*), Ubiquitin Specific Peptidase 9, Y-Linked (*USP9Y*), Eukaryotic Translation Initiation Factor 1A, Y-Linked (*EIF1AY*), which were all downregulated; and X Inactive Specific Transcript (*XIST*), which was upregulated in women with RCC. To determine whether this was a trend for RCC patients, we also compared women with LCC to men with LCC (Table 6). The same genes were altered in women compared to men with RCC versus women and men with LCC. All genes identified as differentially expressed between men and women were located on either X or Y chromosomes and their differential expression frequently occurred in X and Y pairs; i.e. Zinc Finger Protein, X-Linked (*ZFX*) and *ZF* Y-Linked (*ZFY*), *RPS4X* and *RPS4Y1, EIF1AX* and *EIF1AY*.

**Table 5.**
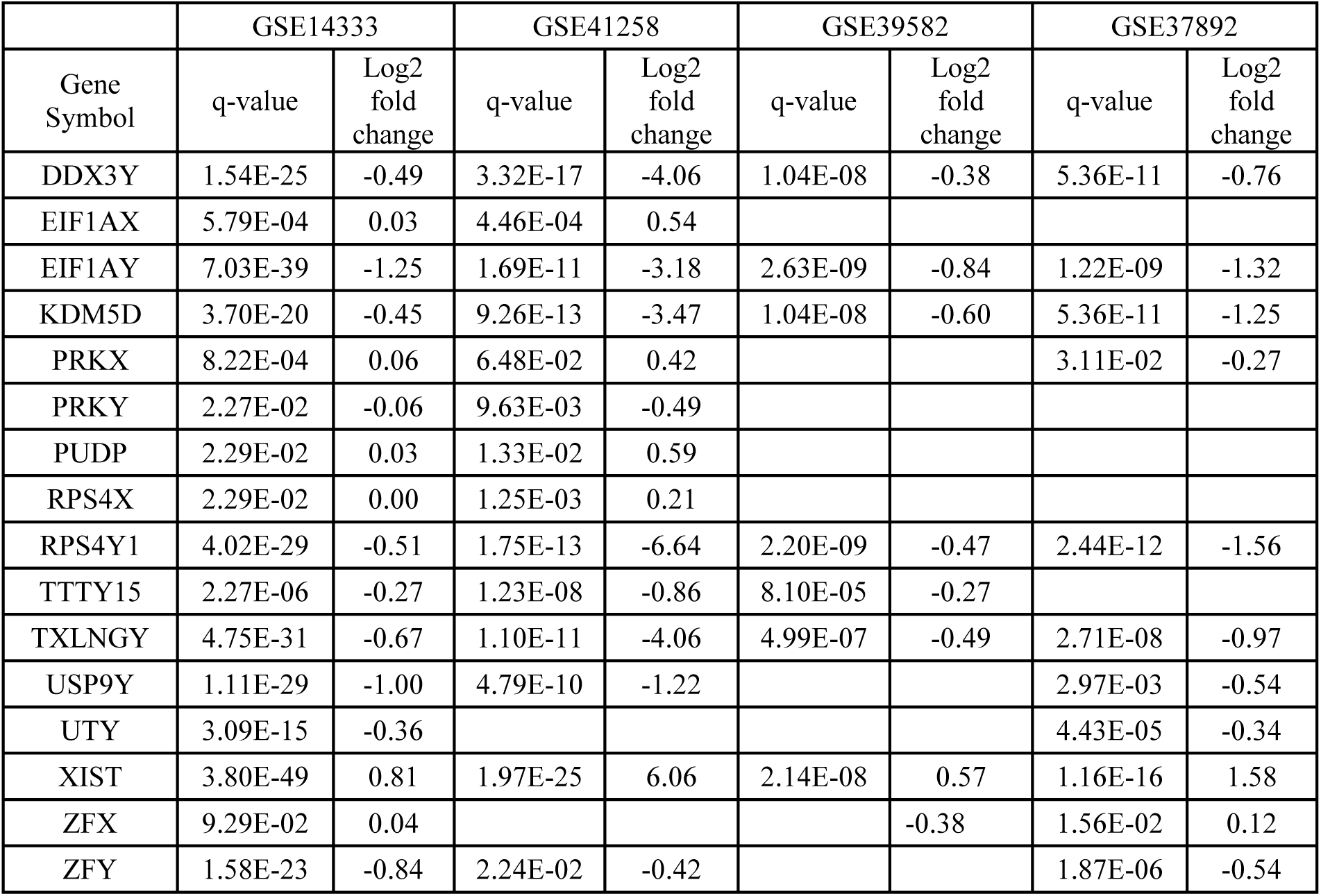
Differential expression of genes in colon tumor from women with RCC compared to men with RCC that reproducibly occur in two or more datasets. Log2 fold changes displayed are with respect to women with RCC.

**Table 6.**
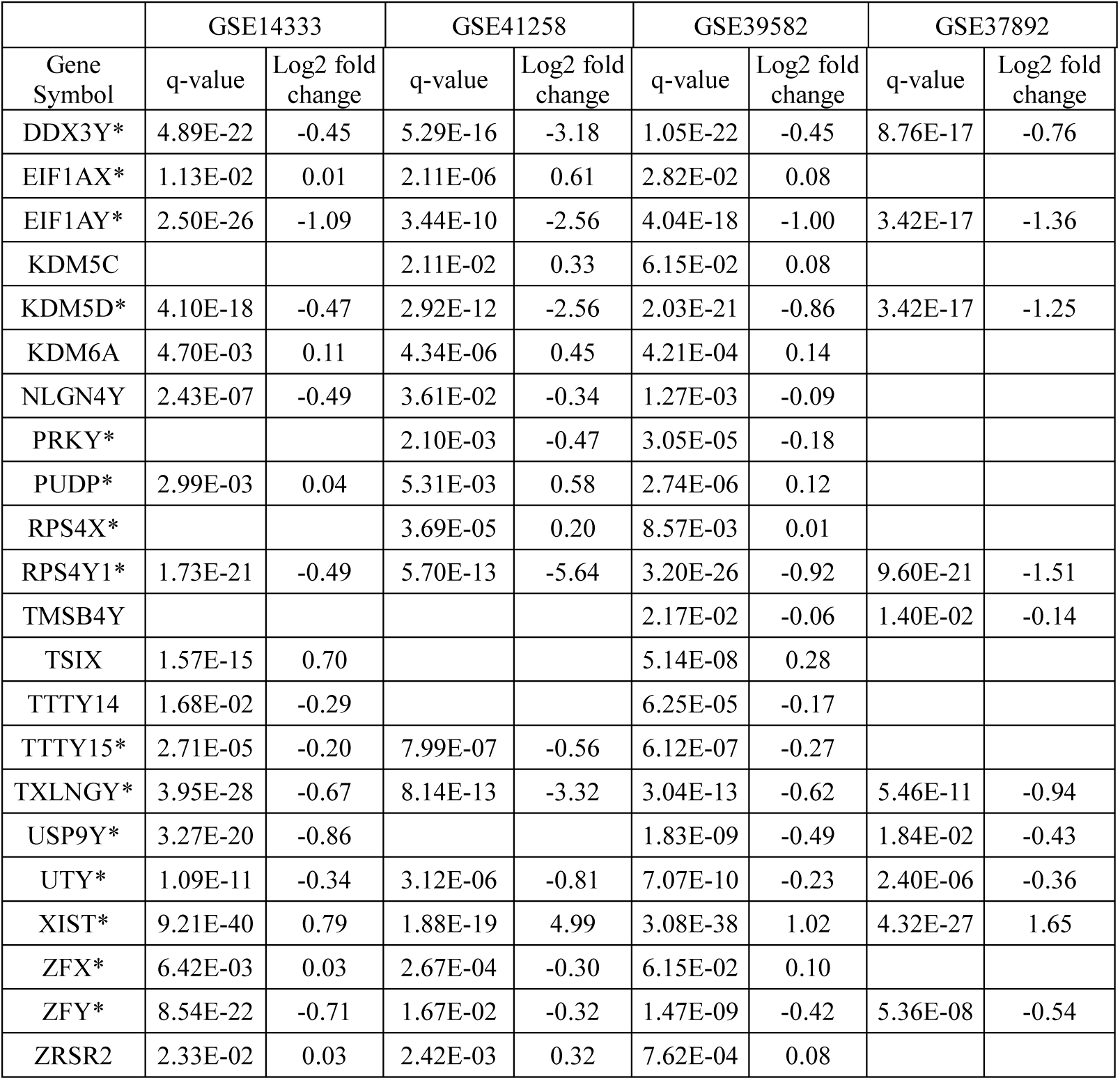
Differential expression of genes in colon tumors from women with LCC compared to men with LCC across four datasets, that reproducibly occur in two or more datasets. Fold changes displayed are with respect to women with LCC. * Genes that are also differentially regulated in women with RCC when compared to men with RCC.

To understand how the lists of differentially expressed genes are linked to the biology and mechanisms of tumor growth and patient survival in women and men with RCC and LCC, we performed enrichment analysis in MetaCore™ software to examine their connectivity in molecular pathways. We also utilized IPA software to identify genetic interplay.

### Enrichment analysis of patients with RCC compared to LCC

Enrichment analysis of differentially expressed genes in women with RCC compared to women with LCC revealed six enriched molecular pathways (FDR <0.05) as illustrated in Figure 2A. We did not identify differentially expressed genes in men with RCC compared to men with LCC with enriched pathway analysis, possibly due to the low number of differentially expressed genes initially identified.

**Figure 2A.**
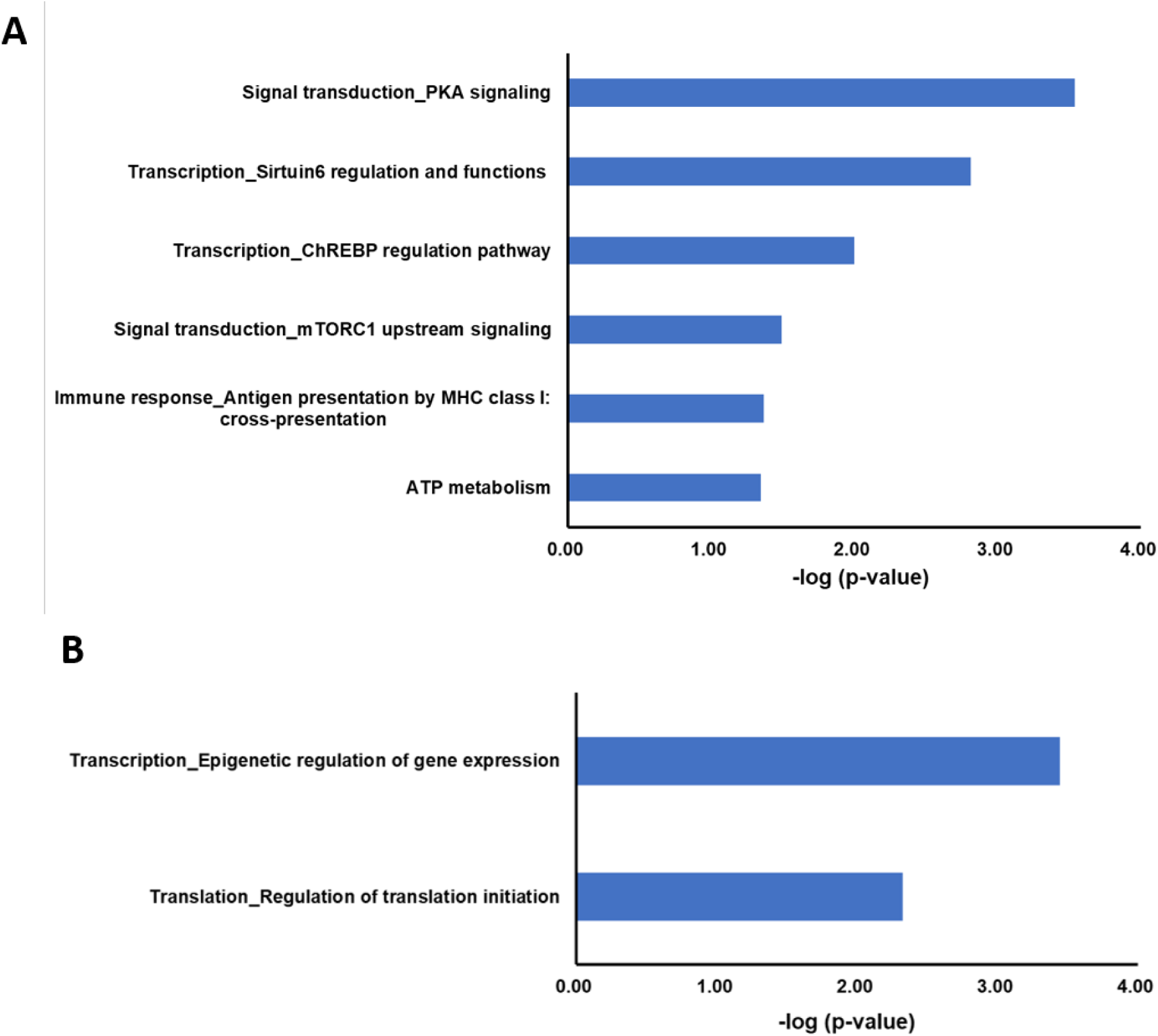
Significantly altered pathways in enrichment analysis when comparing between women with RCC and to women with LCC. **2B.** Significantly altered pathways in enrichment analysis when comparing women with RCC to men with RCC, −log (p-value).

The top significant pathway enriched in women with RCC was the protein kinase A (PKA) pathway (Figure 3). This role of PKA is to phosphorylate and regulate protein activity. PKA is a holoenzyme complex composed of catalytic (PKA-cat) and regulatory (PKA-reg) subunits. PKA-regs exist as two forms: type I (PKA-reg) and type II (PKA-reg type II). When cyclic adenosine monophosphate (cAMP) is bound to PKA-regs, their affinity to PKA-cat is lowered. The PKA holoenzyme thus dissociates and releases PKA-cat to carry out protein phosphorylation. We observed that *PKA-reg* and *PKA-reg type II* expression were downregulated in women with RCC compared to women with LCC causing a potential activation of PKA-cat. This was also supported by the decrease seen in protein kinase inhibitor alpha (*PKI*) which when active inhibits PKA. However, an upregulation in Meprin A subunit beta, which is an inhibitor of PKA catalysis, could affect its activity. Interestingly, increased expression of this gene has been associated with increased cell migration and invasion; thus, its upregulation supports the poorer outcomes seen in RCCs (Wang et al., 2016). Additional genes that were changed in the datasets included cGMP-inhibited 3’-5’-cyclic phosphodiesterase A (*PDE3A*), which was downregulated. Phosphodiesterases regulate cAMP levels through hydrolysis to produce AMP, therefore downregulation of *PDE3A* indicates decreased cleavage of the phosphodiester bond in cAMP. Increased cAMP levels have been shown to be protective against colon cancer (Tsukahara et al., 2013). As cAMP negatively regulates *PKA-reg* this further supports the downregulation of this gene. Protein phosphatase 2 (*PP2A*) expression was also increased, which can lead to increased cell survival in RCC through 5-hydrotryptamine receptor 1A signaling and may have possible actions on the androgen receptor (Dai et al., 2017).

**Figure 3.**
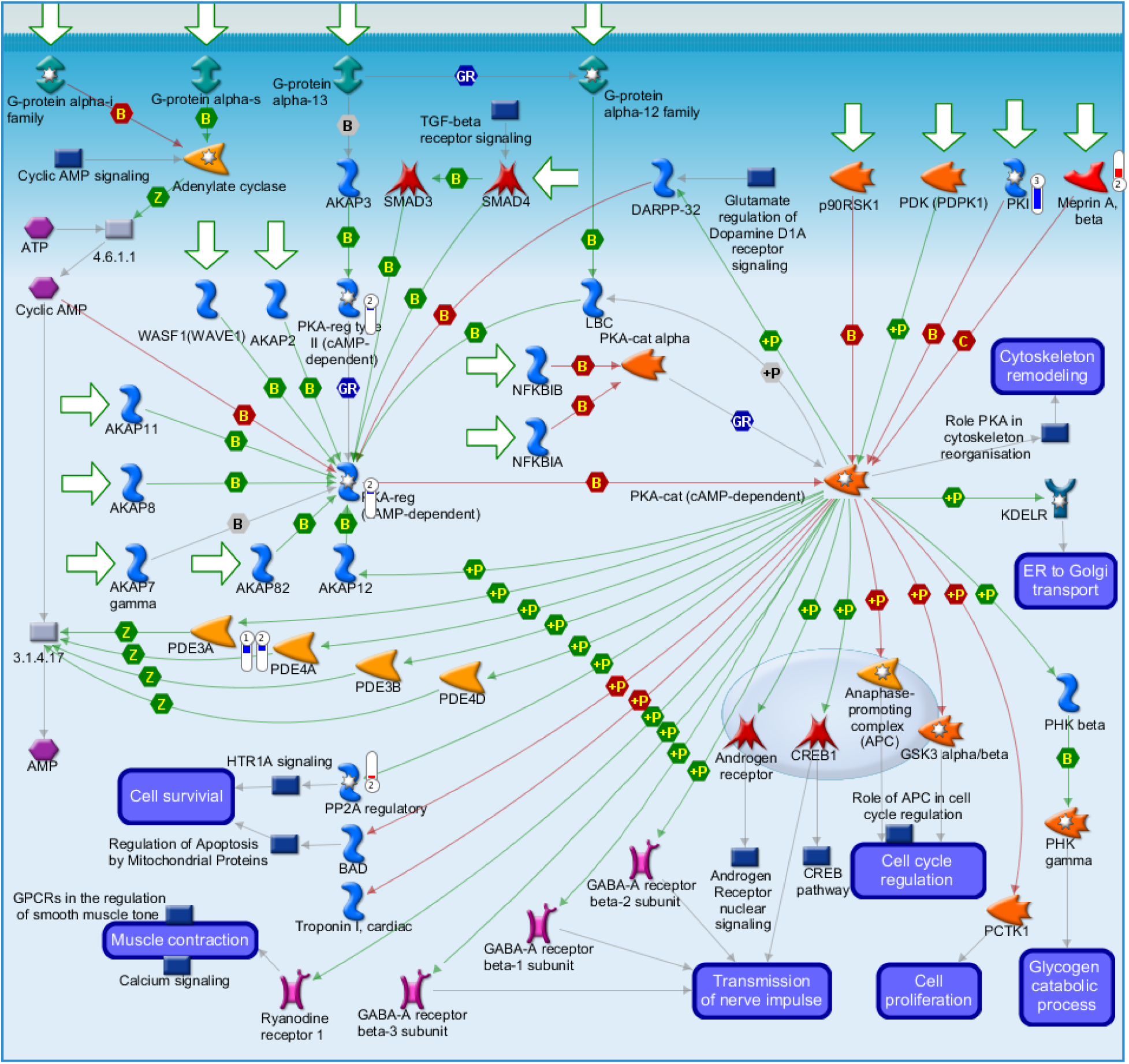
MetaCore generated pathway of differentially expressed genes involved in protein kinase A (PKA) signaling. Experimental data from all three GSE datasets is linked to and visualized on the maps as thermometer-like figures. Upward thermometers have red color and indicate upregulated signals and downward (blue) ones indicate downregulated expression levels of the genes. Annotation are listed in supplemental materials.

The sirtuin (SIRT) 6 pathway was also significantly enriched in women with RCC (Figure 4). One of the roles of sirtuin 6 is to promote an increase in the AMP/ATP ratio, thus regulating energy metabolism in the cancer cell and important metabolic processes in the cell. In our analysis, we observed that 5’ AMP-activated protein kinase alpha subunit (*AMPKα*) was significantly downregulated, which plays a key role in controlling the AMP/ATP ratio. In addition, expression of acyl-coenzyme A oxidase 1 (*ACOX1*), glucokinase (*HXK4*), and Indian hedgehog (*LHH*) genes, all regulated by SIRT6, were decreased. These genes control the synthesis of macromolecules required for cell growth through glucose and fatty acids catabolism and have roles in cellular senescence. Forkhead box O3 (*FOXO3A*) was downregulated, and STIP1 homology and U-box containing protein 1 (*CHIP*) was upregulated, which also have roles in decreasing sirtuin 6 expression and indicates decreased activation of *FOXO3A* by SIRT1. Interestingly, SIRT1 has been implicated in the regulation of cancer cell proliferation through regulation of sex steroid hormones (Moore et al., 2012).

**Figure 4.**
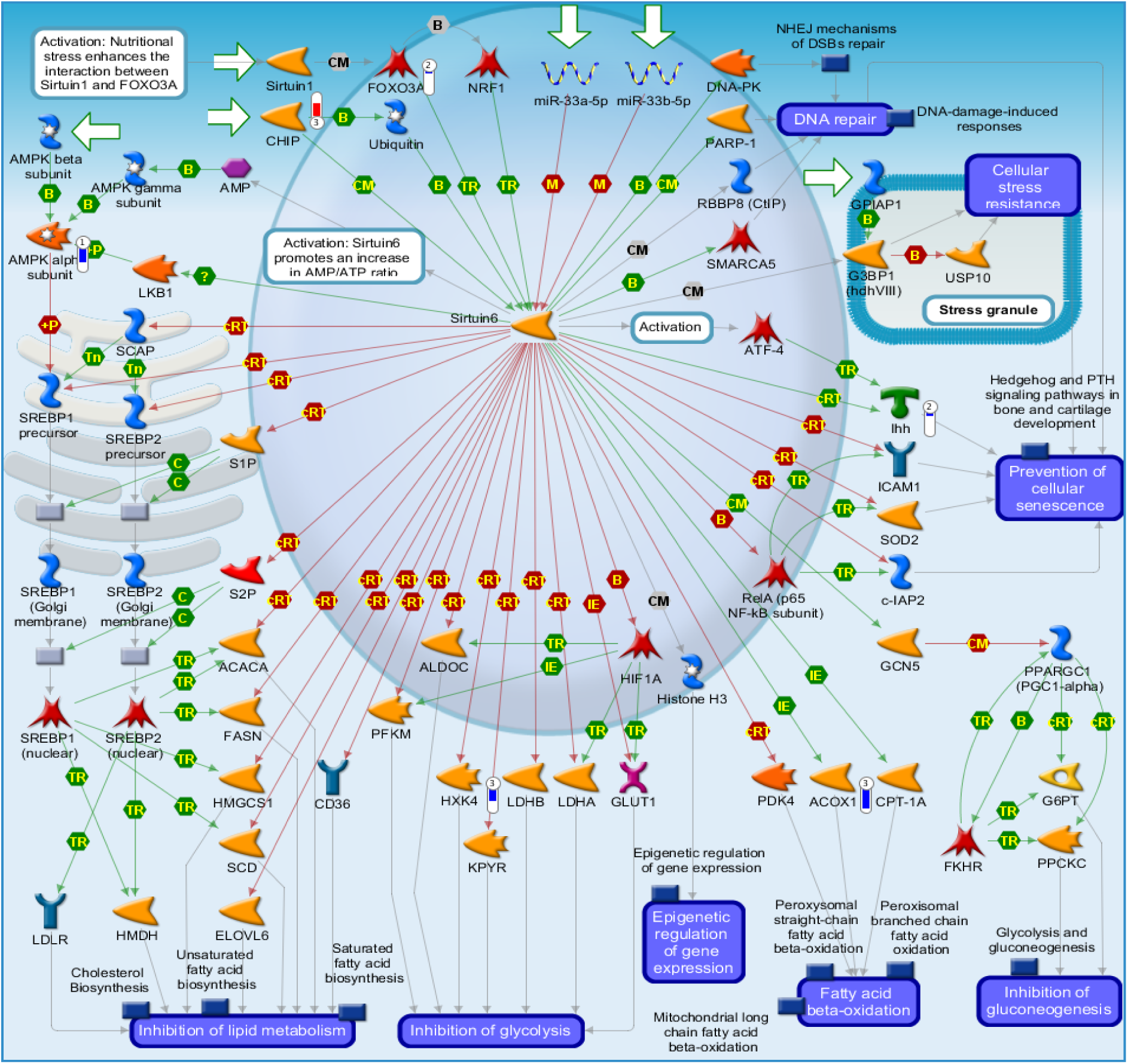
MetaCore-generated pathway showing differentially expressed genes involved in sirtuin 6 regulation and function. Experimental data from all three GSE datasets are linked to and visualized on the maps as thermometer-like figures. Up-ward thermometers have red color and indicate up-regulated signals and down-ward (blue) ones indicate down-regulated expression levels of the genes. Annotation are listed in supplemental materials.

The enrichment of the carbohydrate-responsive element-binding protein (ChREBP) pathway points again to changes in nutrient supply. ChREBP is inhibited by cAMP and PKA, here we observed that *PKA-cat* is activated (due to downregulation of *PKA-reg*), which would decrease phosphorylation and activation of ChREBP. We also observed that *AMPKα* is significantly downregulated along with acyl-coenzyme A synthetase (*ACS*) (Figure 5), which would conversely cause ChREBP activation. Deregulation of these genes as a response to nutrient supply suggests a role for ChREBP-mediated glucose and fatty acid metabolic control and may play a role in cell proliferation. The mammalian target of rapamycin complex 1 (*mTORC1*) signaling pathway was also enriched (Figure 6) suggesting a nutrient deplete environment in women with RCC. Tubulin tyrosine ligase 1 (*TTL1*) and *AMPKα* were both downregulated in women with RCC. mTOR regulation is known to play a role in colon cancer biology (Kimmelman and White, 2017).

**Figure 5.**
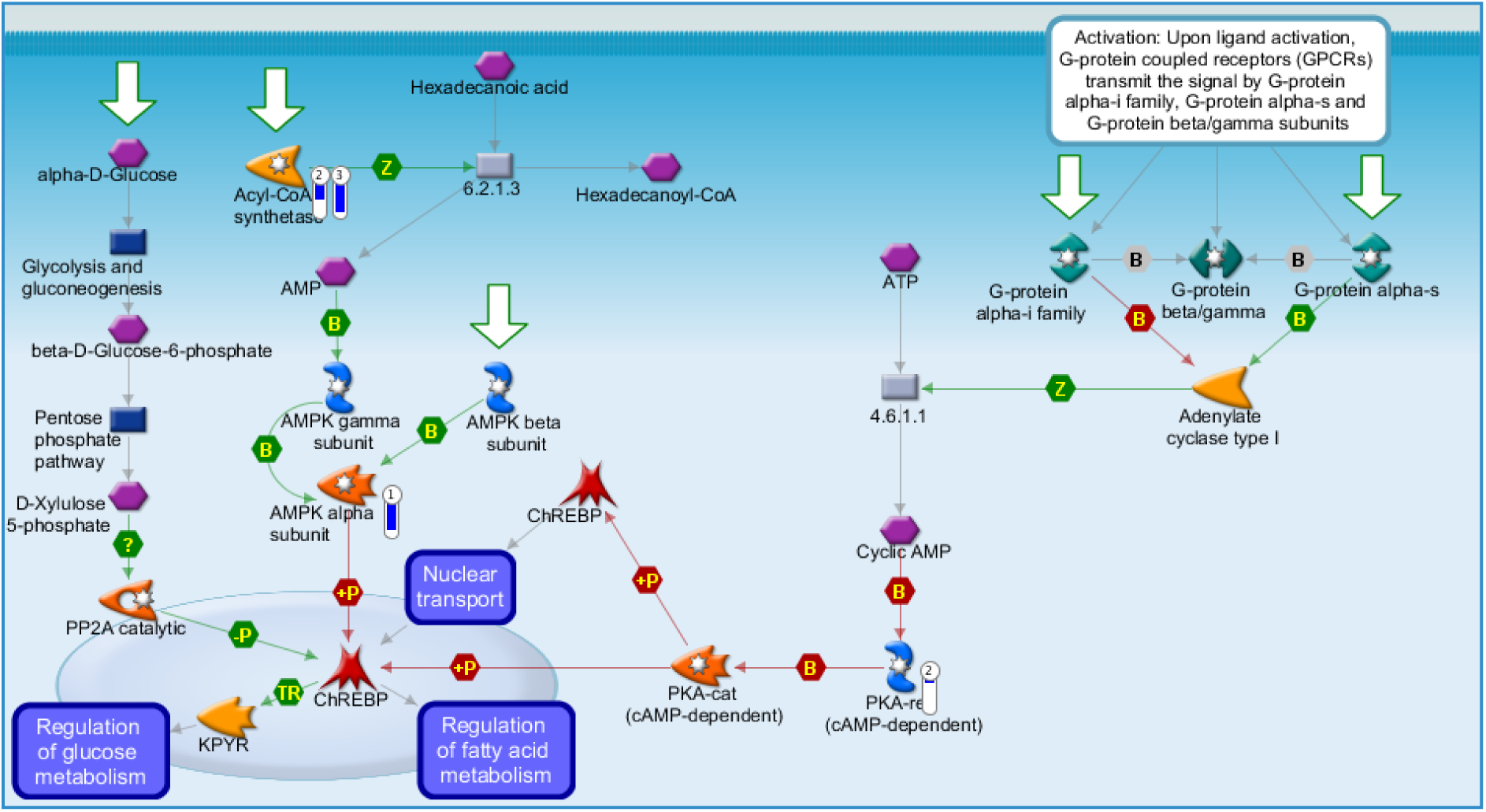
MetaCore-generated pathway showing differentially expressed genes involved in carbohydrate-responsive element-binding protein (ChREBP) regulation. Experimental data from all three GSE datasets are linked to and visualized on the maps as thermometer-like figures. Up-ward thermometers have red color and indicate up-regulated signals and down-ward (blue) ones indicate down-regulated expression levels of the genes. Annotation are listed in supplemental materials.

**Figure 6.**
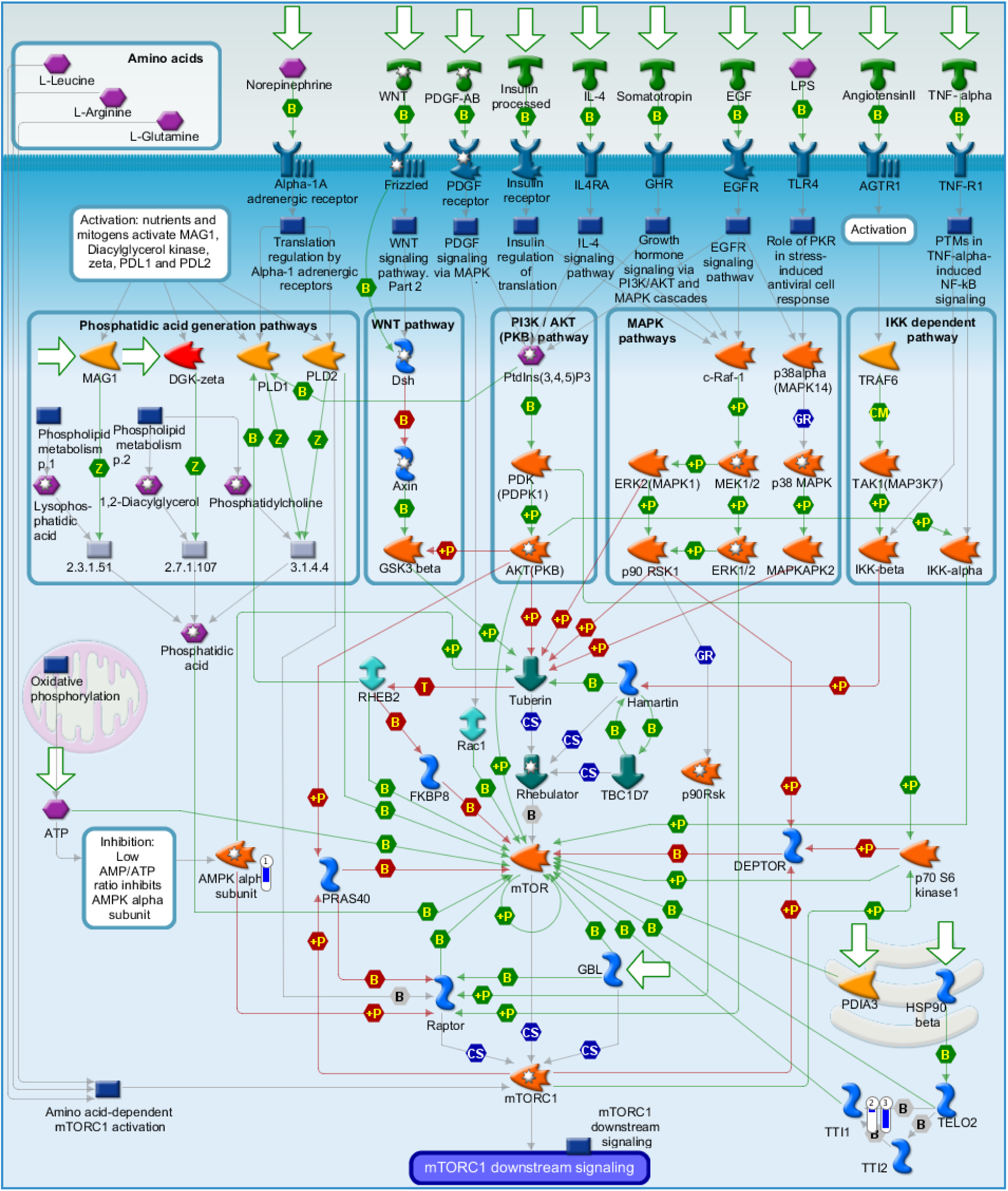
MetaCore-generated pathway showing differentially expressed genes involved in mammalian target of rapamycin complex 1 (mTORC1) regulation. Experimental data from all three GSE datasets are linked to and visualized on the maps as thermometer-like figures. Up-ward thermometers have red color and indicate up-regulated signals and down-ward (blue) ones indicate down-regulated expression levels of the genes. Annotation are listed in supplemental materials.

The ATP metabolic pathway was found to be significantly altered (Figure 7). *PDE3A* and ectonucleotide pyrophosphatase/phosphodiesterase 1 (*ENPP1*) were downregulated, whereas pyruvate kinase muscle isozyme M2 (*PKM2*) and *PDE10A* were upregulated. Downregulation of ENPP1 and PDE3A indicates decreased breakdown of ATP and cAMP respectively, to generate AMP, also increased PKM2 indicates increased ATP production from the metabolism of phosphoenolpyruvate and ADP to pyruvate. Therefore, this pathway also substantiates widespread disruption to AMP and ATP generation.

**Figure 7.**
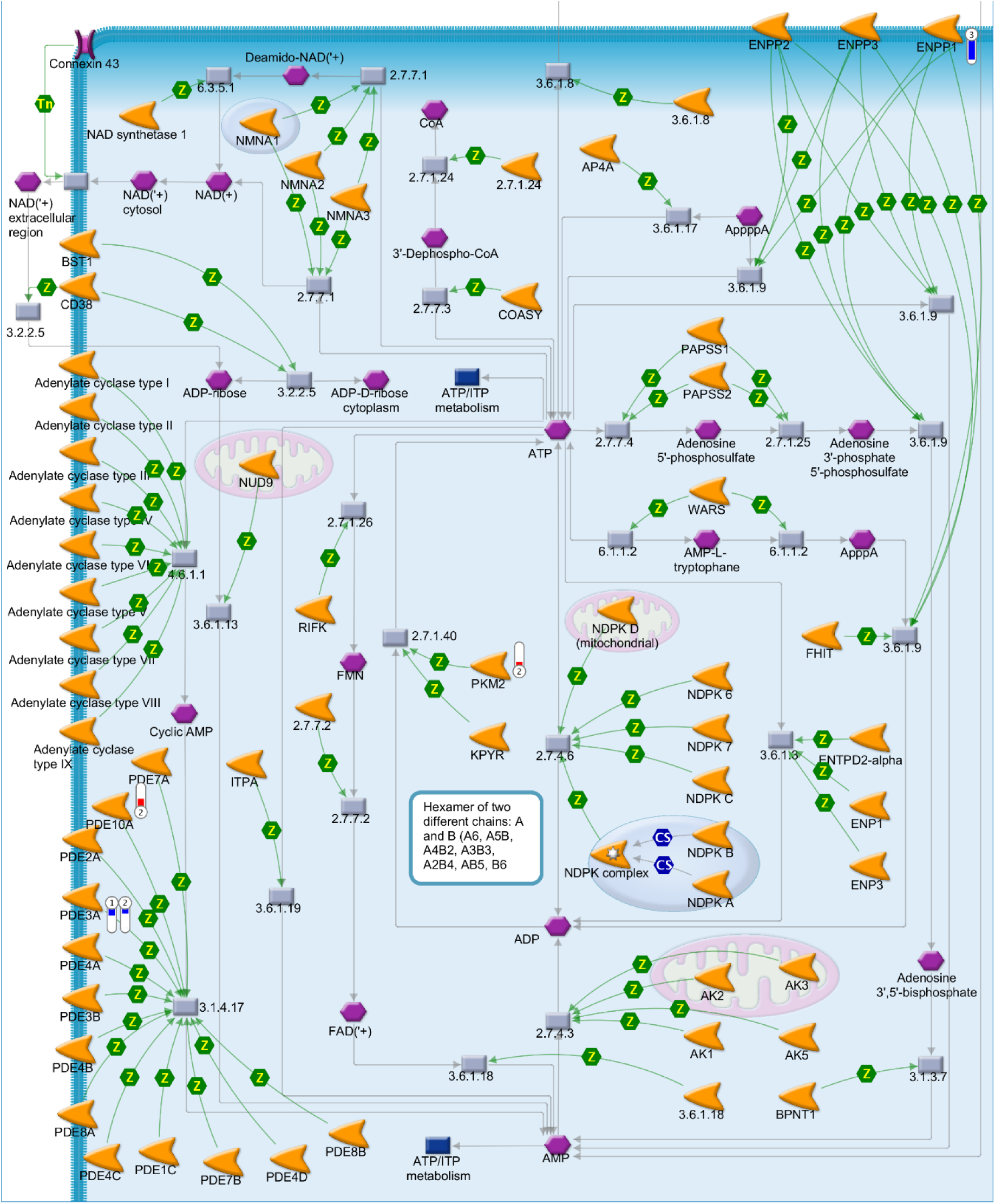
MetaCore-generated pathway showing differentially expressed genes involved in ATP metabolism. Experimental data from all three GSE datasets are linked to and visualized on the maps as thermometer-like figures. Up-ward thermometers have red color and indicate up-regulated signals and down-ward (blue) ones indicate down-regulated expression levels of the genes. Annotation are listed in supplemental materials.

One other distinct mechanism showing significant enrichment in RCC compared to LCCs in women was antigen presentation by major histocompatibility complex class I (Figure 8). All the genes from our datasets that were linked through this pathway were upregulated, suggesting an increase in activity in this pathway; heat shock protein (*HSP*)*70*, hypoxia up-regulated protein 1 (*HYOU1*), and *CHIP* partake in antigen endocytosis, antigen presentation, and T-cell immune response. Of note, the protein encoded by *HYOU1* belongs to the *HSP70* family, which has been shown to be related to cell growth and cancer progression (Jagadish et al., 2016). Interestingly, *HSP105*, a member of the *HSP70* family that has been shown to play a role in anti-tumor immune response, was downregulated (Miyazaki et al., 2005; Yokomine et al., 2006; Yokomine et al., 2007).

**Figure 8.**
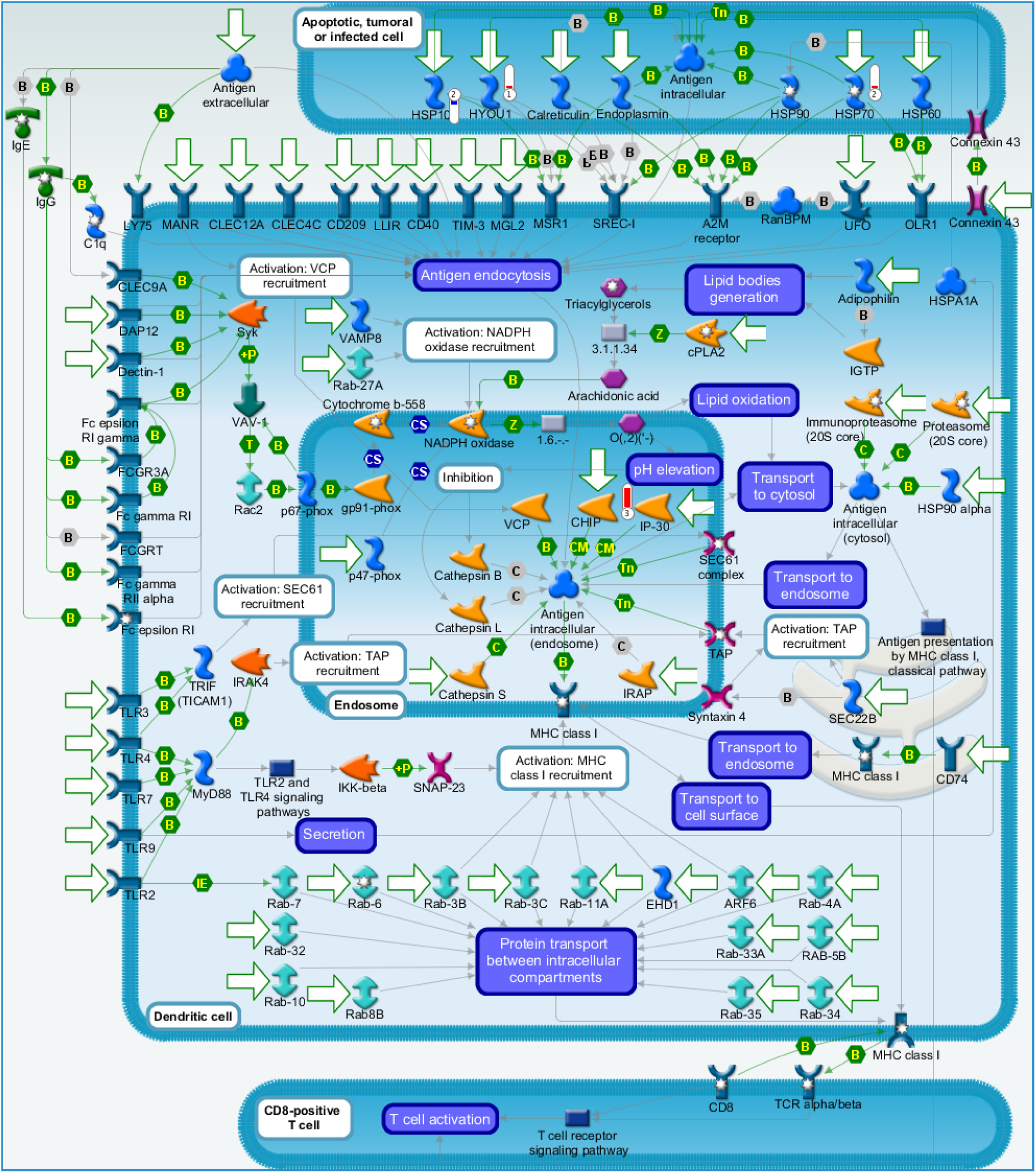
MetaCore-generated pathway differentially expressed genes involved in Antigen presentation by major histocompatibility complex (MHC) class I: cross-presentation. Experimental data from all three GSE datasets are linked to and visualized on the maps as thermometer-like figures. Up-ward thermometers have red color and indicate up-regulated signals and down-ward (blue) ones indicate down-regulated expression levels of the genes. Annotation are listed in supplemental materials.

We also used IPA network analysis to identify significant differentially expressed gene pathways between with women RCC and LCC and revealed possible differences in the regulation of signaling pathways related to cancer cell death and apoptosis (Figure 9). Gene expression differences included the downregulation of protein *O*-fucosyltransferase 1 (*POFUT1*), which is a key factor in the Notch1 (*NOTCH1*) signaling pathway. This pathway is an important regulator for cell death and has been associated with poorer prognosis (Chabanais et al., 2018). Notch 1 known to be essential for maintenance of normal intestinal epithelium and is activated in primary colorectal cancer (CRC) rather than metastatic colon cancer, it may therefore may be more important for early CRC development (Suman et al., 2014). It has also been shown that AMPK depletion can reduce Notch 1 levels (Mohini and Rangarajan, 2018). Although Notch1 is associated with poorer prognosis, our study may imply that Notch1 is not an important factor in RCC versus LCC outcomes. The myelocytomatosis viral oncogene homolog (MYC) and MYC/MAX heterodimer, which play a role in apoptosis, were also downregulated suggesting inhibition of apoptosis. Suppression of MYC has been associated with oxygen and glucose-deprived conditions, and could be a route of cancer cell survival under nutrient depletion (Okuyama et al., 2010). Caspase 6 (*CASP6)*, a protease that plays an important role in apoptosis, was also found to be downregulated in women with RCC. Other involved pathways include the p38 MAPK signaling pathway, a regulator of cell metabolism, proliferation, and invasion/inflammation, and aryl hydrocarbon receptor signaling pathways, which has shown to be involved in tumorigenesis. Thus, clear differences in the expression of genes related to cell growth were seen between RCC and LCC cases in women that could be related to differences in nutrient supply.

**Figure 9.**
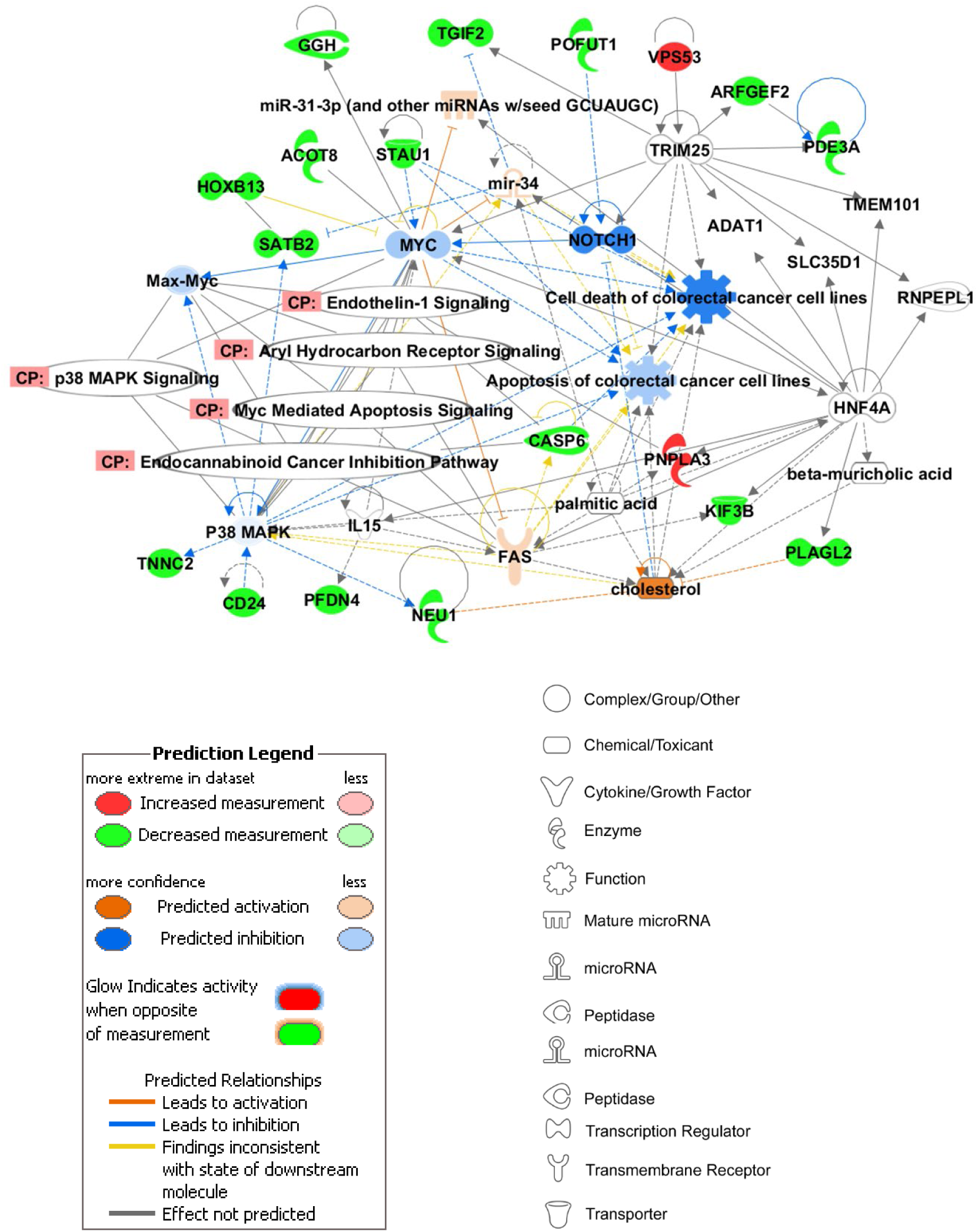
Pathway analysis by Ingenuity Pathway Analysis (IPA). Top IPA network “Cell Morphology, Cell Death, and Survival, Cancer” (Fisher Exact Test p < 1×10E-48) generated from the differentially expressed genes between women with RCC vs LCC (Table 3). Molecules colored in green are downregulated, red-colored are upregulated. Molecules in white are added by the IPA network generating algorithm to complete the network.

### Enrichment analysis of patients with RCC comparing men to women

There were two significantly enriched pathways seen in all datasets from women with RCC when comparing to men with RCC and were related to expression of genes involved in regulation of transcription and translation (Figure 2B). When comparing men and women with LCC, we saw similar enrichments of these pathways (Transcription_Epigenetic regulation of gene expression; p-value (-log) 3.44, Translation_Regulation of translation initiation; p-value (-log) 2.33). The genes that underlie the variation in expression between men and women are located on either chromosome X or Y, thus it is not surprising that differences between these genes are identified when comparing men to women. An example of this can be seen when comparing the *EIF1A* genes. *EIF1AY* is upregulated in men when comparing differences between the two sexes in either RCC or LCC patients. However, most of the genes identified have links to cancer and reveal the importance of their increased or decreased expression in this disease.

## Discussion

Identification of differentially expressed genes and pathways in RCC and LCC is a potentially powerful means of prognostication and could provide predictive markers for response to treatment of colon cancer. In this study, we use multiple publicly available gene expression datasets to examine gene expression in tumors of right and left sided colon cancer in men and women and identify statistically significant and clinically relevant differences in signaling pathways. When comparing women with RCC to women with LCC, there were four genes that were commonly dysregulated across two or more large datasets and had fold changes >2; *ARID3A, SATB2*, and *TNNC2* that were downregulated in women with RCC, and *HOXC6* that was upregulated.

*ARID3A* codes for a DNA-binding protein and is proposed to be a tumor suppressor; higher expression of *ARID3A* is correlated with increased overall survival and correlated with p53 status (Song et al., 2014a; Song et al., 2014b). In addition, high *ARID3A* expression is more frequently observed in microsatellite-stable (MSS) and microsatellite-instable (MSI)-low cases versus MSI-high cases (MSI-high is more often observed in RCC) (Song et al., 2014a). *SATB2* is a transcription factor that regulates chromatin remodeling and transcription (Mansour et al., 2015). High expression of *SATB2* and *ARID3A* were recently shown to be a biomarker for favorable prognosis and sensitivity to chemotherapy and radiation of colon cancer and potential metastasis (Zhang et al., 2018). We found downregulation of these genes in women with RCC compared to women with LCC, which may play a role in poorer prognosis of this patient group. Limited publications have yet to discuss the implications of decreased expression of *TNNC2* in RCCs. However, higher expression has been shown to correlate with decreased survival (the human protein atlas/TCGA data).

*HOXC6,* a transcription factor belonging to the family of human homeobox (*HOX*) genes that control cell morphogenesis and differentiation during embryological development, known to be expressed in differential gradients to establish craniocaudal (head to tail) polarization. *HOXC6* is also known to be overexpressed in numerous cancer types. Specifically, it is associated with poor survival in colon cancer patients. Upregulation of this gene, which was found to be upregulated in our study in both men and women with RCC, has been shown to correlate with poor overall survival in right sided colon cancer, and is thought to promote carcinogenesis via inhibition of autophagy and mTOR pathway activation (Ji et al., 2016a; Ji et al., 2016b). We saw upregulation in men and women with RCC compared to LCC, which is in concordance with prior studies. Interestingly, *HOXC6* modulates androgen receptor (AR)-stimulated gene expression, and thus, could play a role in hormonal mechanisms in cancer (Ramachandran et al., 2005).

Gene expression profiles from men with RCC and LCC also revealed differences in gene expression between the two sides of the colon. However, the fold changes and number of genes observed were less than those seen in women. *HOXC6* was upregulated with fold change >2 in women with RCC. *MUC12* and *PRAC1* were similarly downregulated in both men and women when comparing RCCs to LCCs. Recent studies have examined mRNA expression using data from TCGA and GEO comparing patients with RCC to LCC have revealed similar changes to *HOXC6, PRAC1* and *MUC12* (Hu et al., 2018; Peng et al., 2018). The *PRAC1* gene is associated with hypermethylation, and reported to be expressed in the prostate, rectum and left-sided colon. It is also downregulated in patients with prostate cancer and in immortalized cell lines from RCC patients (Bauer et al., 2012). Downregulation of *PRAC1* is thought to be repressed *via* hypermethylation of one of the differentially methylated regions situated on the CpG island shore in RCCs. It is also hypothesized to be a tumor suppressor gene through its interactions with cotranscribed *HOXB13* (downregulated in women with RCC when compared to women with LCC in our analysis, but not differentially expressed in men) (Hu et al., 2018). *MUC12* is one of the *MUC* genes that code for glycoproteins important for mucosal barrier function, decreased *MUC12* expression has been correlated with poorer survival for stage II and III CRC patients (Matsuyama et al., 2010).

On direct comparison of tumors from women with RCC to men with RCC, fold changes of >2 were observed in gene expression. However, the genes observed were specific to sex-chromosomal location (X or Y-linked). Thus, the expression of a Y-chromosomal linked genes would be expected to be higher in men when comparing to women.

Pathway analysis was carried out to identify the relationships between the genes expressed and their involvement in metabolic pathways to determine biological processes related to right or left-sided colon cancer in men and women. Only enrichment analysis of differentially expressed genes from women revealed significantly enriched pathways with respect to tumor location in the colon. The six pathways which are the most highly enriched in relationship to occurrence of cancer in RCC in women involve signaling, metabolism, and immune response. Five of the pathways observed are involved in the regulation of essential nutrients such as glucose and fatty acids, and control the generation of energy metabolites such as AMP and ATP. There is a clear link between high ATP and cAMP, low AMP, accompanied by a decrease in *AMPKα* expression in women with RCC. ATP is involved in processes that mediate all types of cell death including apoptosis, autophagy and necrosis, thus plays a critical role in the survival of cancer cells (Zhou et al., 2012). AMPK*α* is involved in the regulation of multiple metabolic functions in the cell, and when activated, it can stimulate glycolysis, inhibit fatty acid synthesis and promote fatty acid oxidation under conditions of nutrient depletion. When AMPK*α* is induced under these conditions, it also plays a role in autophagy via suppression of mTORC1 and activation of unc-51-like autophagy activating kinase 1 (ULK1) (Inoki et al., 2003). Conversely, pathway analysis also revealed a potential downregulation of mTORC1 signaling in women with RCC through decreased regulation by *TTL1* expression (Figure 6). In addition, gene expression analysis revealed an increase in *HOXC6* expression, which has been shown to be linked to this pathway in CRC cells via the promotion of autophagy and inhibition of mTOR (Ji et al., 2016a). One of the main roles of mTORC1 is to sense nutrient availability (primarily amino acids, cellular energy (via AMPK), and oxygen levels) to control cell growth. When mTORC1 is inactivated it dissociates from the ULK1 complex which in turn activates ULK1. Activation of ULK1 is essential for autophagy and the involvement of AMPK*α* in this process is also required (Rabanal-Ruiz et al., 2017). However, it has been shown that under nutrient deplete conditions, ULK1 can directly phosphorylate and downregulate AMPK at the *α* subunit, thus providing a negative regulatory feedback loop decreasing autophagy (Loffler et al., 2011), which may explain the differences we observe within this pathway. An additional examination of *ULK1* and autophagy (*ATG*) gene expression in the four datasets did not show significant differences between women with RCC compared to women with LCC, however analysis of mTORC1/ULK1 phosphorylation and signaling would help to identify the association of autophagy to women with RCC. *FOXO3A* was also downregulated and highlighted in the SIRT6 pathway. FOXO3 has been implicated in the transcriptional regulation of autophagy and functions in parallel with the mTOR pathway. However, unlike mTOR, autophagy by FOXO3 is dependent on the transcriptional upregulation of multiple autophagy genes such as *ATG12, ATG4B, VPS34, ULK2, LC3B, GABARAPL1, BECLIN1, BNIP3*, and *BNIP3L* those that were measured in the datasets (the latter six) were not significantly dysregulated.

Therefore, due to the potential therapeutic targeting of autophagy in cancer, which can promote cancer progression, the survival of tumors under stress conditions, and response to chemotherapeutics, the association of autophagy in women with RCC is worth further investigation (Mokarram et al., 2017).

The other distinct pathway which was enriched in women with RCC compared to LCCs was immune regulation. This finding is also in agreement with the recent assignment of tumor subtypes that are associated with sex and colon location (Guinney et al., 2015; Lee et al., 2017a). Consensus Molecular Subtype (CMS1) tumors, those with high immune infiltration and activation, were more frequently diagnosed in women with RCC (where their definition of RCC including transverse colon). Whereas CMS3, classified as a metabolically active subtype, does not appear to be more prevalent in one side of the colon than the other. However, their classification of LCC included rectum which may influence the difference seen in our findings. Additionally, we found upregulation of genes encoding heat shock proteins *HSP70* and downregulation of *HSP105* in women with RCC. These proteins are also involved in immune response. Upregulation of *HSP70* expression has been shown to increase cell proliferation and tumor growth, and has been associated with poorer prognosis in colon cancer (Jagadish et al., 2016). *HSP105* partakes in increasing anti-tumor immune response, which was downregulated in RCC (Miyazaki et al., 2005; Jagadish et al., 2016). Our findings of the dysregulation of these heat shock proteins in women with RCC versus LCC may play a role in the more aggressive nature and generally poorer prognosis of RCC.

Therefore, this study provides a comprehensive bioinformatic analysis of differentially expressed genes and pathways, commonly altered among different sexes and anatomical locations in colon cancer. It also shows potential therapeutic targets for treatment involving suppressors and activators in altered pathways. The results lead us to the overall hypothesis that women with RCC have inactivation of *AMPKα*, and a decreased AMP/ATP ratio in their tumor tissues when compared to women with LCC. This study also highlights the importance and value of open science by using publicly available datasets to provide novel findings and prove reproducibility in our findings between datasets.

## Supporting information

Supplemental Key

## Acknowledgements

The authors would like to thank Women’s Health Research at Yale, and the Yale Cancer Center for funding this project (CJ, SK, YZ). This publication was also made possible by CTSA Grant Number UL1 TR001863 from the National Center for Advancing Translational Science (NCATS), components of the National Institutes of Health (NIH), and NIH roadmap for Medical Research (SK). Its contents are solely the responsibility of the authors and do not necessarily represent the official view of NIH.

## Author Contribution Statement

CHJ conceptualized the study, YS, EPFL, RGM, analyzed the data. YS, VM, YC, QZ, YC, YZ, VV, SAK, CHJ analyzed the results and helped write the publication. YS, VM, YC, RGM, CHJ helped edit the manuscript and prepare for publication.

