## Supplementary figures and images for "Molecular pathway analysis indicates a distinct metabolic and immune phenotype in women with right-sided colon cancer"

### Supplemental Key

**Key for understanding MetaCore-generated figures**

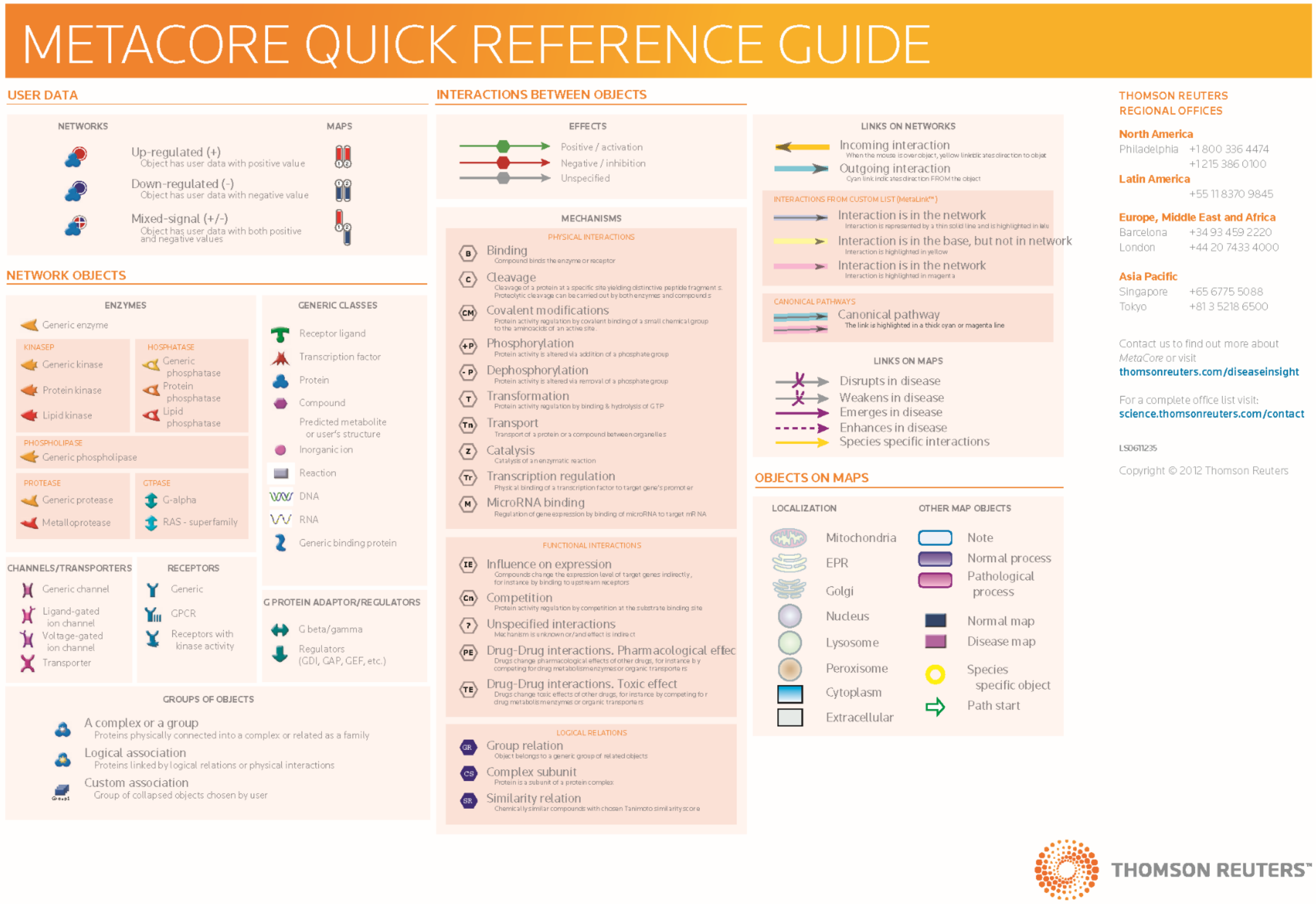
